# Signaling and transcriptional dynamics underlying early adaptation to oncogenic BRAF inhibition

**DOI:** 10.1101/2024.02.19.581004

**Authors:** Cameron T. Flower, Chunmei Liu, Hui-Yu Chuang, Xiaoyang Ye, Hanjun Cheng, James R. Heath, Wei Wei, Forest M. White

**Author notes:** These authors contributed equally to this study (Cameron T. Flower, Chunmei Liu).

## Abstract

A major contributor to poor sensitivity to anti-cancer kinase inhibitor therapy is drug-induced cellular adaptation, whereby remodeling of signaling and gene regulatory networks permits a drug-tolerant phenotype. Here, we resolve the scale and kinetics of critical subcellular events following oncogenic kinase inhibition and preceding cell cycle re-entry, using mass spectrometry-based phosphoproteomics and RNA sequencing to capture molecular snapshots within the first minutes, hours, and days of BRAF kinase inhibitor exposure in a human *BRAF*-mutant melanoma model of adaptive therapy resistance. By enriching specific phospho-motifs associated with mitogenic kinase activity, we monitored the dynamics of thousands of growth- and survival-related protein phosphorylation events under oncogenic BRAF inhibition and drug removal. We observed early and sustained inhibition of the BRAF-ERK axis, gradual downregulation of canonical cell cycle-dependent signals, and three distinct and reversible phase transitions toward quiescence. Statistical inference of kinetically-defined signaling and transcriptional modules revealed a concerted response to oncogenic BRAF inhibition and a dominant compensatory induction of SRC family kinase (SFK) signaling, which we found to be at least partially driven by accumulation of reactive oxygen species via impaired redox homeostasis. This induction sensitized cells to co-treatment with an SFK inhibitor across a panel of patient-derived melanoma cell lines and in an orthotopic mouse xenograft model, underscoring the translational potential for measuring the early temporal dynamics of signaling and transcriptional networks under therapeutic challenge.

## Introduction

Cellular adaptation to exogenous sources of stress is driven by coordinated changes in cell state, often involving interactions between hundreds of distinct genes and proteins over diverse timescales. In the case of oncoprotein inhibition in cancer by targeted therapy treatment, such responses can promote rapid adaptation to drug, enabling survival of tumor cells and treatment failure.^1,2,3,4^ A well-studied example of this phenomenon is the response to receptor tyrosine kinase (RTK)-RAS-ERK pathway inhibitors in certain cancers, where decreased activity of the target kinase can potentiate other mitogenic signaling proteins due to reduced negative-regulatory feedback.^5,6^ Notably, such adaptive processes often unfold on timescales that are inconsistent with selection of (epi)genetically resistant subclones, but can influence eventual selection by defining new heritable cell states or promoting endogenous hypermutagenic processes, thereby promoting long-term acquired resistance.^7,8^ Understanding early adaptive responses to therapy can yield promising new combination therapies which block compensatory signaling axes, preventing tumor cells from persisting and driving disease progression.^9^

Examinations of the signaling responses to targeted therapy have yielded valuable insights into the biochemical drivers of drug tolerance and cancer cell persistence. However, many efforts to date have focused on interrogating particular pathways of interest based on prior knowledge, further emphasizing these pathways’ importance in drug adaptation but leaving most adaptive signaling mechanisms underexplored. In principle, adaptation to exogenous perturbation may involve the remodeling of multiple complex signaling networks over unpredictable timescales; capturing such network remodeling requires methods for time-resolved quantitation of systems-level signaling networks.^10,11^ While large-scale ‘hypothesis-free’ efforts have gained traction in recent years with the falling costs of next-generation sequencing and mass spectrometry (MS)-based proteomics, the dynamics associated with early adaptation to targeted therapy have remained poorly resolved, as most reported hypothesis-free studies to date have profiled with low temporal granularity under drug treatment or have focused on the events following cell cycle re-entry. Here, we performed a time-resolved analysis of early adaptive signaling using a patient-derived cell line of BRAF^V600E^-mutant melanoma, which is characterized by an incomplete response to BRAF inhibitor therapy and therefore serves as a prototypical model system to chart the early molecular events that permit cancer cell survival following oncogenic signaling blockade. Prior work has established the propensity of these cells to undergo substantial transcriptional, epigenetic, and metabolic remodeling under continued BRAF inhibition, promoting survival and eventual slow re-cycling under drug.^12,13,14,15,16^ However, the links between these events and loss of BRAF kinase activity, as well as the intricacies of how cell-wide signaling networks contribute to the onset of drug adaptation within the initial days of exposure, remain unclear. To address this knowledge gap, we used quantitative MS to monitor the temporal dynamics of thousands of protein phosphorylation sites over a three-day period of BRAF inhibition, followed by a six-day drug washout, to assemble and explore a highly detailed map of cellular signaling kinetics under targeted therapy treatment.

## Results

### Profiling signaling network dynamics in a patient-derived melanoma model of adaptive BRAFi resistance

To examine the early adaptive response to targeted therapy exposure, we studied a patient-derived melanoma cell line, M397, that has previously been shown to exhibit tolerance to BRAF inhibition despite harboring the BRAF^V600E^ oncogenic mutation.^12,13,14,15,16^ We confirmed M397 cells exhibit a biphasic viability response to the BRAF inhibitor vemurafenib (VEM) after 72 hours; within the first response phase, the concentration of VEM necessary to attain half-maximal viability (EC50, 98.37 ± 5.41 nM, mean ± standard deviation) was within range of the reported IC50 of VEM for BRAF (32 nM and 100 nM against mutant- and wild-type BRAF, respectively^17^), consistent with an on-target response (Figure 1A). Significantly higher doses of VEM (>30 μM) drove cell death presumably through substantial off-target activity, and significantly lower doses (<10 nM) induced a hormetic effect that has previously been linked to paradoxical activation of BRAF.^18^ At a dose centered between the two response phases (3 μM), cells fail to apoptose and instead show a growth-arrested phenotype beginning at 30 hours (Figures 1B and S1A), indicating the suitability of M397 as a model system for studying the mechanisms of survival and early adaptation to oncogenic kinase inhibition.

**Figure 1.**
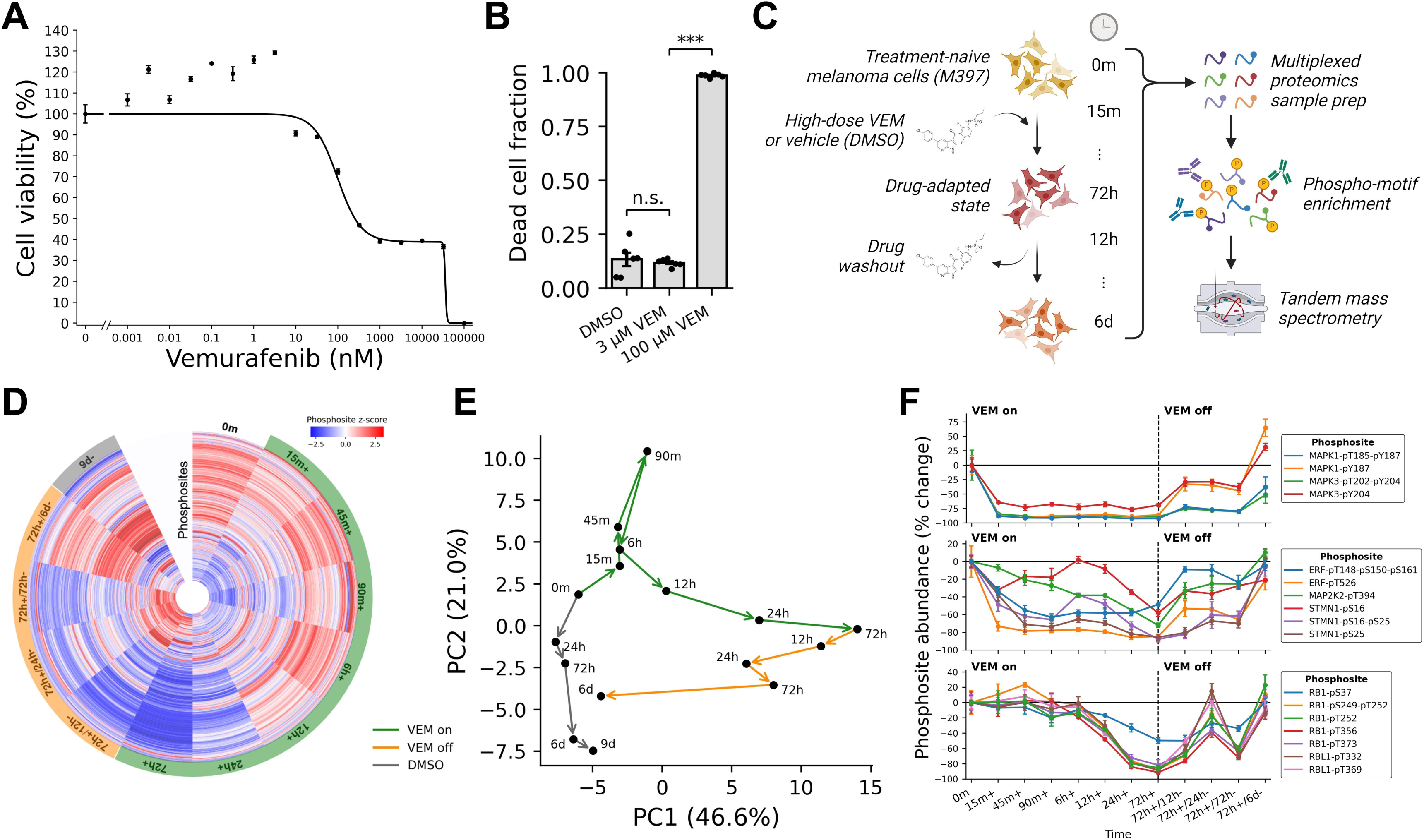
Early signaling dynamics of oncogenic BRAF inhibition in melanoma cells. A. Viability dose-response of a patient-derived BRAF-mutant melanoma cell line, M397, against BRAF inhibitor vemurafenib (VEM). Error bars depict standard error of the mean across three replicates. B. Fraction of dead cells (staining positive for trypan blue) under high doses of VEM. Measurements were collected 72 hours after drug treatment. Error bars depict standard error of the mean across three replicates. *** : *P* < 0.005 by two-sided *t*-test. C. Time-resolved phosphoproteomics workflow schematic. D. Protein phosphorylation dynamics under BRAF inhibition by high-dose VEM (3 μM). Abundance values were derived from the mean across three biological replicates followed by z-score normalization. Phosphosites with no change throughout the time course (<10% coefficient of variation across timepoints, 108 total) are absent from this visual. E. Principal components analysis (PCA) of phosphoproteome dynamics. F. Phosphosite dynamics of MAPK1 and MAPK3 (ERK2 and ERK1, respectively) (top), canonical ERK1/2 substrates (middle), and RB1/L1 (bottom). Error bars depict standard error of the mean across three replicates.

Treatment of *BRAF*-mutant cells with high-dose VEM is expected to rapidly ablate the BRAF-ERK axis and drive widespread remodeling of subcellular signaling networks. To examine this response, we treated M397 cells with VEM at 3 μM in biological triplicate and harvested from culture at seven early timepoints, spanning 15 minutes to 72 hours (Figure 1C and Supplementary Table S1). A separate group of cells were treated with 0.1% (v/v) of drug solvent dimethyl sulfoxide (DMSO) and harvested at a subset of the timepoints. After cells had been exposed to VEM for 72 hours, drugged media was replaced with drug-free media for six additional days; this drug washout period allowed us to assess the reversibility of adaptation. All lysates were then subjected to multiplexed quantitative phosphoproteomics by MS using tandem mass tags (see Methods). We chose to prioritize particular kinase families known to mediate survival, cycling, and adaptation to stress including tyrosine kinases, DNA damage-sensing kinases, mitogen-activated protein kinases (MAPKs), and cyclin-dependent kinases (CDKs). To increase the probability of detection and accurate quantitation of the substrates of these kinases by MS, their corresponding phospho-motifs were serially immunoprecipitated from the pool of TMT-labeled tryptic peptides prior to MS analysis (Figure S1B). A total of 4,160 phosphopeptides covering 4,088 phosphorylation sites were confidently identified and quantified, and 1,461 sites were detected in all three independent replicates (1,273 sites when counting multi-phosphorylated and non-unique peptide mappings as single sites).

The time-resolved phosphoproteome showed a rapid response to BRAF inhibition within 15 minutes followed by waves of distinct signaling network remodeling and recovery (Figure 1D). Principal components analysis (PCA) of the phosphoproteomics data revealed an early (0 to 90 minutes) response to VEM along principal component 2, followed by a delayed (90 minutes to 72 hours) response along principal component 1 which was almost fully reversed following six days of drug removal (Figure 1E). Under drug washout, cells traverse a distinct path through principal component space which terminates near the matched DMSO control samples, indicating the controls successfully captured the signaling changes associated with sustained culture and low-level DMSO exposure. Pairwise correlation analysis followed by hierarchical clustering between all samples showed strong consistency between biological replicates and revealed three distinct timescales of drug response: ‘early’ (15 to 90 minutes), ‘intermediate’ (6 to 12 hours), and ‘late’ (24 to 72 hours) (Figure S1C). Notably, all drug washout samples co-cluster with ‘late’ timepoints, with the exception of the final washout timepoint (72h+/6d-BR1-3) which co-clusters with ‘basal’ samples, indicating a heavily delayed but nevertheless reversible signaling response.

The ERK MAPKs, MAPK1 (ERK2) and MAPK3 (ERK1), which are canonically activated downstream of active BRAF, were rapidly and durably inhibited by VEM treatment throughout the time course until drug removal, as measured by activation loop phosphorylation (Figures 1F and S1D, top) and by phosphorylation of canonical ERK1/2 substrates (Figures 1F and S1D, middle). The inhibition of ERK1/2 within 15 minutes of VEM exposure confirms successful drug-target engagement, and the durability of inhibition suggests M397 cells are not dependent on reactivation of the RAF-ERK axis to survive high-dose VEM treatment, in contrast to observations of rapid ERK1/2 reactivation in some cancers treated with RTK-RAS-ERK pathway inhibitors.^5,6,19^

Activation of CDKs, especially CDK4/6, canonically occurs downstream of ERK1/2 activation and promotes entry into S-phase in part by hyperphosphorylation of RB1.^20^ We observed gradual downregulation of seven phosphosites on RB1 and the related protein RBL1 under VEM, and recovery under drug washout, consistent with reversible cell cycle arrest at G1-phase (Figures 1F and S1D, bottom). These findings confirm that our molecular snapshots capture drug-tolerant cells in a period of transient, reversible stress driven by oncogenic kinase inhibition and further support the M397 model’s utility for scrutinizing the dynamics of early drug adaptation.

### The time-resolved transcriptome informs dynamic gene regulatory responses to oncogenic BRAF inhibition

Changes in cellular signaling network activity often lead to widespread transcriptional remodeling. To characterize concomitant transcriptomic alterations following signaling network rewiring, we subjected VEM-treated M397 cells to bulk mRNA sequencing (RNA-seq) at select timepoints (Supplementary Table S2). PCA of the time-resolved transcriptome showed strong overall agreement with the phosphoproteome, highlighting immediate-early responses along principal component 1, delayed responses along principal component 2, and a near-complete reversal of the adaptive response during drug washout (Figure 2A). These transitions were characterized by changes in proliferation, metabolism, and stemness along two distinct timescales with a clear transition point at 24 hours. Notably, mRNA transcripts of canonical targets downstream of ERK1/2 signaling showed reduced abundance under VEM and recovery under drug washout, consistent with the durable and reversible ERK1/2 inhibition observed in the phosphoproteome (Figure 2B).

**Figure 2.**
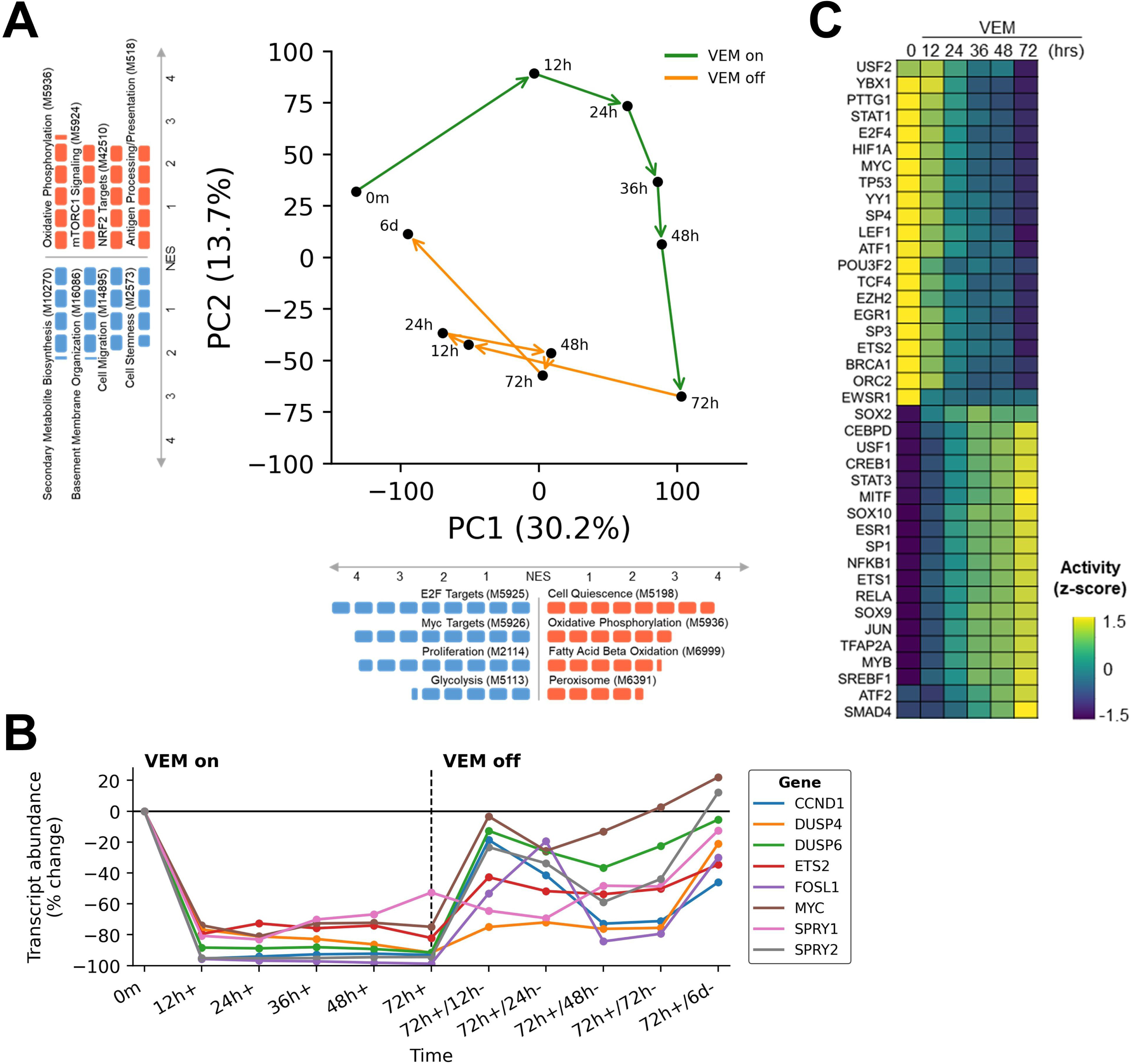
Transcriptional dynamics of oncogenic BRAF inhibition. A. Principal components analysis (PCA) of transcriptome dynamics under high-dose VEM (3 μM), and a subset of significantly enriched gene sets (FDR < 0.05) along each principal component. NES: normalized enrichment score. B. mRNA dynamics of canonical downstream transcriptional targets of the RAS-ERK pathway. C. Inferred transcription factor activity dynamics under VEM.

We performed computational inference of transcription factor (TF) activity to examine the dynamic gene regulatory consequences of oncogenic BRAF inhibition. Estimates of the relative activity of TFs using the time-series RNA-seq data revealed escalating activity of MITF, SOX9, and SOX10 under VEM, alongside diminishing activity of E2F4, HIF1A, MYC, TP53, and ETS2, aligning with an augmented melanocytic signature, attenuated ERK1/2 activity, and cell cycle arrest upon short-term BRAF inhibition (Figure 2C).^12,21^ We further discerned increased TF activities of NFκB and JUN/AP1, corroborating previous reports.^16,22,23^ Construction of a TF regulatory network, using the correlative interactions among TFs whose activities changed significantly within the three days of drug exposure, spotlighted the known mutually activating interactions among the melanocytic lineage TFs MITF, SOX9, and SOX10, in addition to mutually inhibitory interactions between MYC and JUN, and between MYC and NFκB/RelA (Figure S2),^16,22,24,25,26^ further validating our measurements of the early drug-altered transcriptome.

### Integrative analysis reveals the concerted signaling and transcriptional responses to BRAF inhibition

In general, signaling and transcription constitute tightly interconnected processes. Leveraging the time resolution of our multimodal data, we performed an integrative, correlation-based analysis to examine the coordinated network responses to BRAF inhibition. We subjected the phosphoproteomics and transcriptomics data separately to pairwise correlation analysis followed by hierarchical clustering, resulting in data-driven groupings of phosphosites (Figure 3A) and genes (Figure 3B). As each module consists of dozens to hundreds of features which share a kinetically similar response to VEM treatment and removal (Figures S3A and S3B), this analysis served as a form of dimensionality reduction, projecting both datasets into an interpretable kinetically-defined space. Upon examining meta-correlations between inferred signaling and transcriptional modules (Figure S3C), we found our approach detected known ground-truth associations between signaling and transcription; for instance, signaling module A (hereafter, SM-A), which contained phosphosites important for ERK1/2 signaling, was significantly correlated with transcriptional module L (hereafter, TM-L), a growth-enriched module with significant overrepresentation of downstream effectors of canonical RAS-ERK signaling (Figure 3C). SM-A was also significantly anticorrelated with transcription module E (TM-E), which was gradually induced by VEM and overwhelmingly represented genes associated with metabolic rewiring, especially fatty acid and lipid metabolism, consistent with the established role of ERK1/2 in repressing expression of fatty acid β-oxidation genes.^27,28^ Another pair of independently inferred modules, SM-I and TM-J, were downregulated by VEM with near-identical kinetics and were both strongly associated with cell cycle progression and RNA processing (Figure S3D). These findings collectively validate our statistical framework for detecting positive and negative functional links between signaling networks and gene regulation.

**Figure 3.**
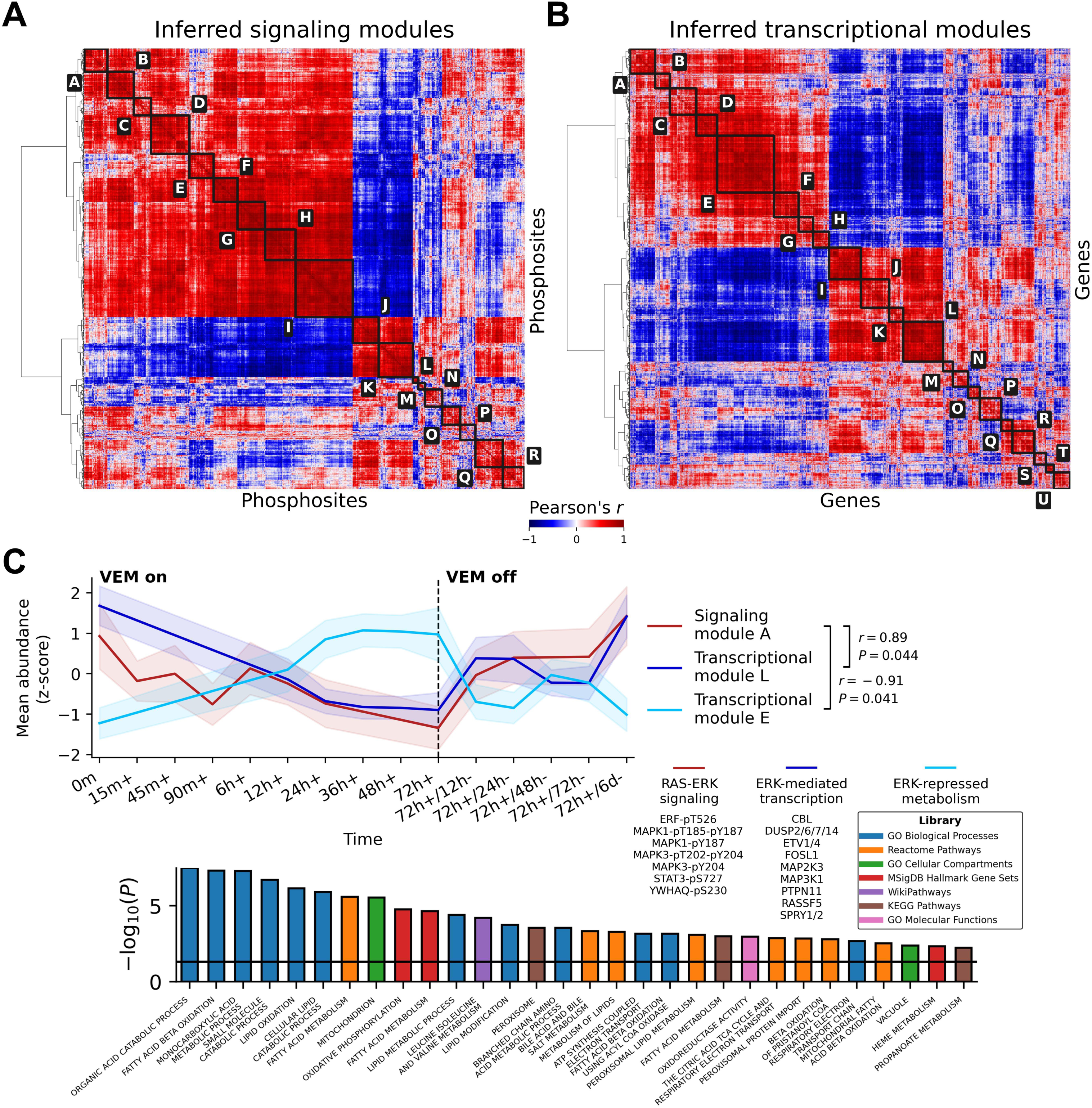
Integrative analysis reveals the concerted molecular response to VEM. A. Inference of kinetically distinct signaling modules from time-series phosphoproteomics data by pairwise correlation analysis and hierarchical clustering. B. Inference of kinetically distinct transcription modules from time-series transcriptomics data by pairwise correlation analysis and hierarchical clustering. C. Consensus dynamics of signaling module I and transcription module J under VEM (top left); phosphosites in signaling module I related to cell cycle progression and RNA processing (top right); overrepresented gene sets in transcription module J related to cell cycle progression and RNA processing (bottom). Shaded regions depict standard deviation across all members of the depicted module. *P*-values were derived from two-sided Fisher’s exact test with multiple hypothesis correction by the Benjamini-Hochberg procedure.

While many of the proteins in the observed phosphoproteome are known to play important functional roles in processes enriched within the correlated transcriptome, most of their phosphorylation sites are poorly characterized. For example, EIF3B is a translation initiation factor that engages the 40S ribosome and is thought to be required for several of the initial steps of protein synthesis, but the precise functional role of the phosphorylation site quantified in SM-I (pS95) is unknown. As the vast majority of the phosphoproteome still lacks confidently annotated functional roles,^29^ this time-resolved map between signaling dynamics and transcriptional remodeling under a well-controlled perturbation may serve as an information-rich resource to nominate mechanistic hypotheses with phosphosite-specific resolution.

### BRAF inhibition induces early tyrosine kinase signaling and cytoskeletal remodeling

While most of the detected protein phosphorylation sites were downregulated by VEM, we found two signaling modules, SM-J and SM-K, which were reversibly induced rather than inhibited by VEM treatment (Figure 4A). Among all phosphosites in these modules, we observed significant depletion of the (pS/pT)P phospho-motif, consistent with reduced MAPK and CDK activity, and significant overrepresentation of phosphotyrosine, suggesting a net increase in tyrosine kinase activity under VEM (Figures 4B and 4C).

**Figure 4.**
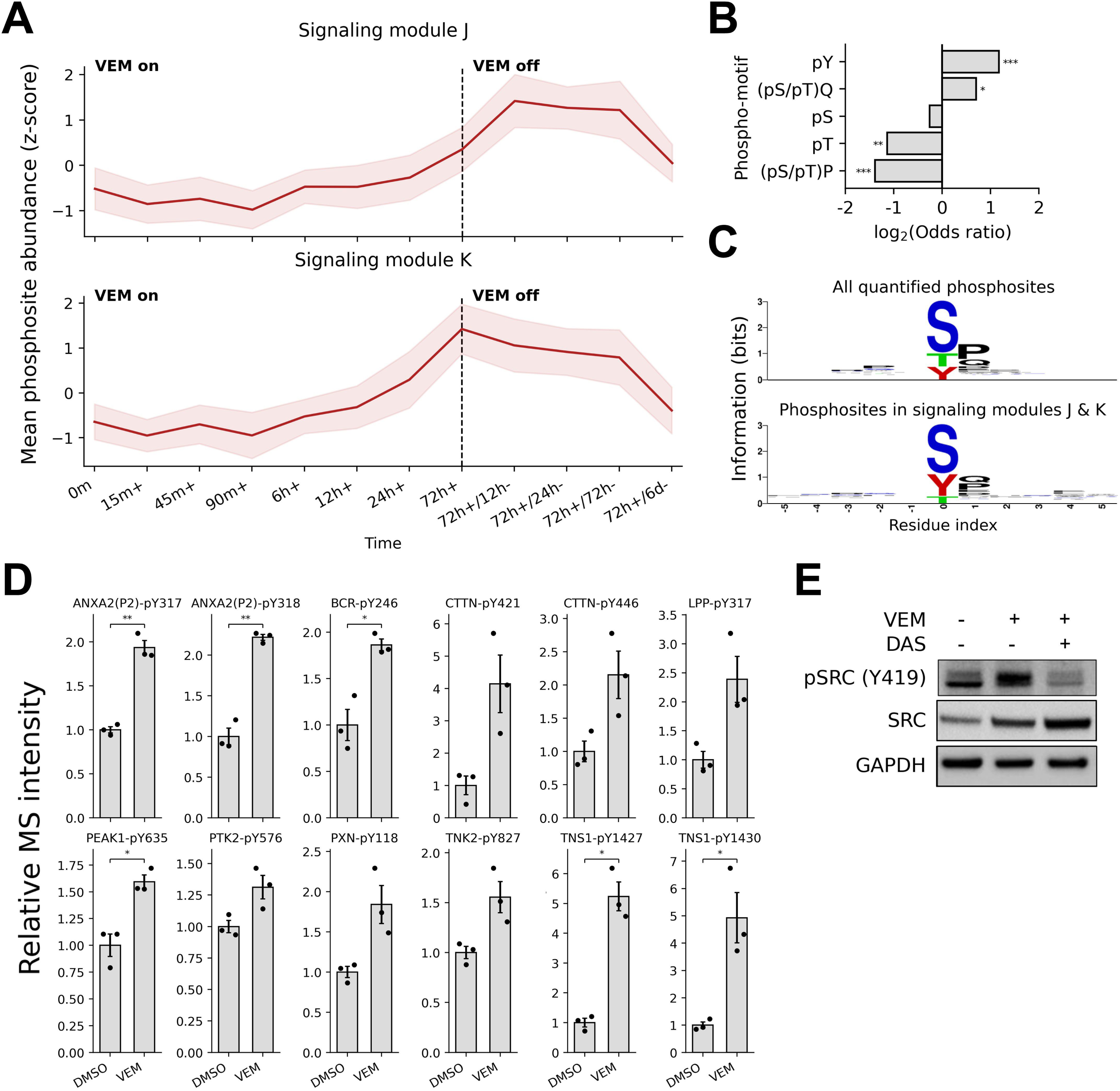
BRAF inhibition leads to an increase in tyrosine phosphorylation, SRC family kinase (SFK) signaling, and cytoskeletal remodeling. A. Consensus dynamics of VEM-induced signaling modules J (top) and K (bottom). Shaded regions depict standard deviation across all phosphosites in the depicted module. B. Representation of each immunoprecipitated phospho-motif among phosphosites in signaling modules J and K. * : *P* < 0.05; ** : *P* < 0.01; *** : *P* < 0.005 by two-sided Fisher’s exact test with multiple hypothesis correction by the Benjamini-Hochberg procedure. C. Consensus sequence logos flanking all phosphosites in the phosphoproteomics data (top) and phosphosites in signaling modules J and K (bottom). Multi-phosphorylated peptides were excluded from the sequence logos. D. Abundances of phosphosites on known substrates of SFKs at 72 hours under VEM or DMSO. Error bars depict standard error of the mean across three replicates. * : *P* < 0.05; ** : *P* < 0.01 by two-sided *t*-test with multiple hypothesis correction by the Benjamini-Hochberg procedure. E. Immunoblot of phosphorylated SRC activation loop and total SRC under VEM alone and in combination with SFK inhibitor dasatinib (DAS).

Upon inspection of the tyrosine-phosphorylated members of SM-J and SM-K, we found that many are known substrates of one or more SRC family kinases (SFKs) (Figure 4D and Supplementary Table S3). SFKs constitute a group of non-receptor tyrosine kinases that coordinate important signaling events regulating cell morphology, survival, and metabolism, among many other biological processes, and have been linked to adaptation and resistance to targeted therapy.^30,31,32,33^ Many of these phosphosites, in addition to other known SFK substrates which were not in SM-J or SM-K, were upregulated under VEM treatment after 72 hours compared to DMSO-treated cells, suggesting their induction was not a consequence of sustained culture. Notably, some of these sites did not show corresponding increases in transcript abundance, indicating that VEM treatment modulates SFK signaling by mechanisms that are at least partly uncoupled from transcriptional upregulation (Figure S4A).

By western blot, we observed increased phosphorylation of the SRC activation loop at Y419 after treatment with VEM (3 μM) for 72 hours, which was prevented by co-treatment with the SFK inhibitor dasatinib (DAS) (Figure 4E). Consistent with the critical role of SFK signaling in regulation of the cytoskeleton and focal adhesions, cells showed marked changes in morphology under VEM treatment (Figure S4B). These results, together with our observation of SFK substrate kinetics, suggest that VEM treatment drives a gradual increase in SFK activity within the first three days of treatment.

### Accumulation of reactive oxygen species under BRAF inhibition promotes SFK signaling

Signaling through SFKs is complex and is known to be driven by many possible cell-intrinsic and extrinsic factors. One known mechanism for SFK activation under cell stress is through accumulation of reactive oxygen species (ROS), such as hydrogen peroxide (H2O2), which can oxidize catalytic cysteine residues in tyrosine kinase domains, resulting in increased kinase activity.^34,35,36^ ROS can also alter cysteine residues on protein tyrosine phosphatases (PTPs), leading to inactivation of PTPs and a net increase in phosphotyrosine signaling.^37,38^ Targeted therapies often promote intracellular accumulation of ROS due to the disruption of redox homeostasis;^39,40^ the resulting oxidative stress can be detrimental to cellular function and thereby contribute to the efficacy of therapy, but when maintained at sub-lethal levels ROS can serve as a second messenger to promote cellular adaptation in the face of external stressors by activating survival-promoting pathways. Within the inferred transcriptional module TM-J, which was significantly anticorrelated with the SFK-associated signaling module SM-K, we found significant overrepresentation of genes associated with NRF2 (NFE2L2), NRF1 (NFE2L1), and maintenance of oxidative stress, suggesting cells had undergone a reduced antioxidant response under VEM (Figure 5A). NRF2 is a TF that promotes cytoprotection from ROS by binding to antioxidant response elements (AREs) in the promoters of target genes including those encoding oxidoreductases and other detoxifying enzymes.^41^ Direct examination of NRF2 transcript levels showed reduced expression under VEM and recovery with VEM removal, and nearly identical transcriptional kinetics of the NRF2 binding partner MAFG (Figure 5B). Additionally, VEM treatment prompted a concomitant gradual increase in expression of the E3 ubiquitin ligase β-TrCP (BTRC), which promotes NRF2 turnover,^42^ and of MAF, a transcriptional repressor that targets AREs for suppression,^43^ throughout the time course. These nearly-synchronized transcriptional events suggest VEM-treated cells undergo gradual dysregulation of redox homeostasis over the first three days of treatment.

**Figure 5.**
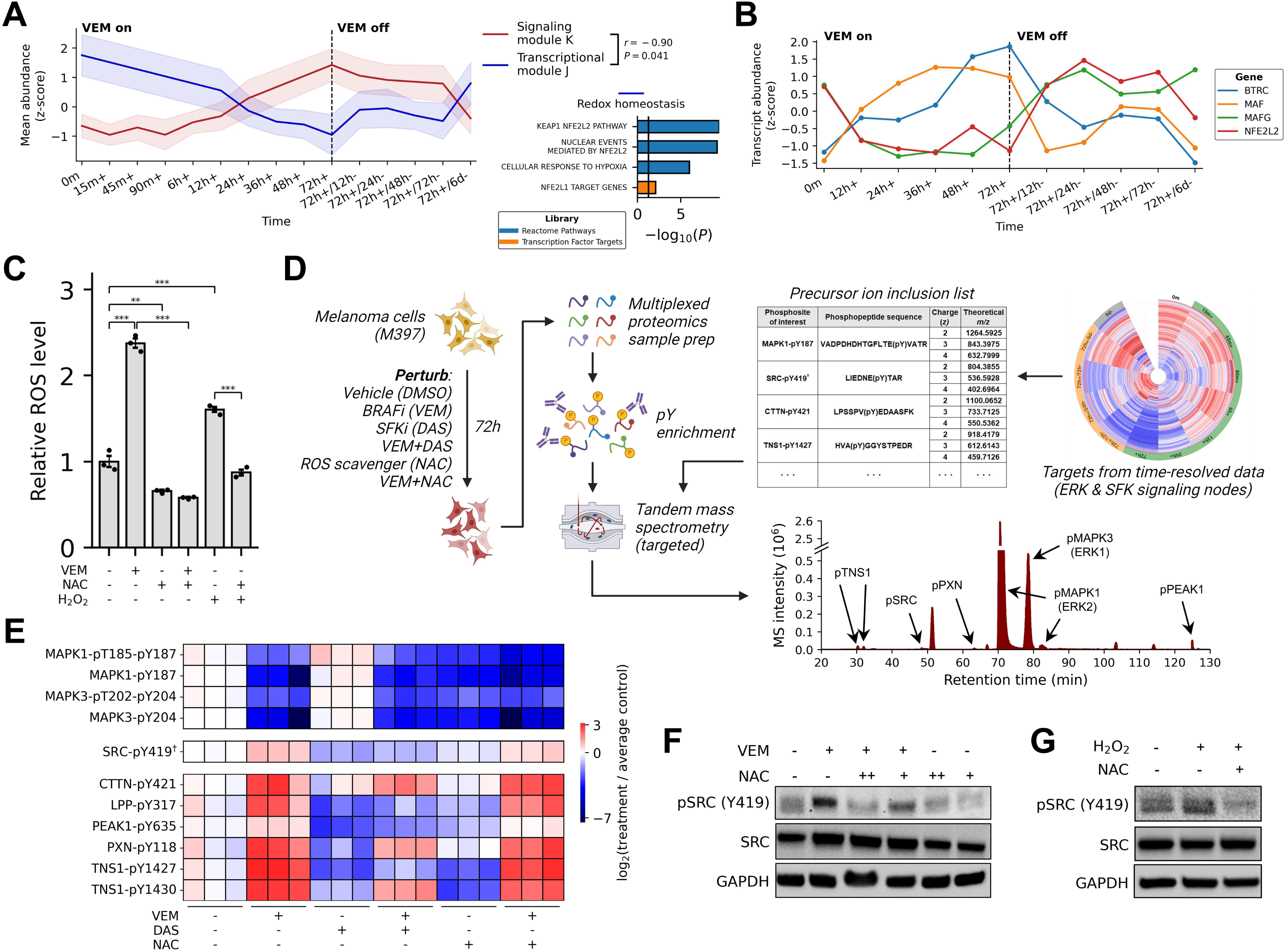
Reactive oxygen species (ROS) accumulation under BRAF inhibition promotes SFK signaling. A. Consensus dynamics of signaling module K (SM-K) and transcription module J (TM-J) (left); overrepresented gene sets in TM-J related to maintenance of oxidative stress (right). *P*-values were derived from two-sided Fisher’s exact test with multiple hypothesis correction by the Benjamini-Hochberg procedure. B. Transcriptional dynamics of antioxidant-promoting transcription factor NRF2 (NFE2L2) and related redox homeostasis genes. C. Abundance of ROS in M397 cells treated with VEM (3 μM), antioxidant prodrug N-acetylcysteine (NAC) (10 mM), hydrogen peroxide (H_2_O_2_) (30 μM), and combinations thereof. All treatments were delivered for 72 hours except NAC, which was delivered one hour before measurement. Error bars depict standard error of the mean across three replicates. ** : *P* < 0.01; *** : *P* < 0.005 by two-sided *t*-test. D. Workflow schematic for targeted mass spectrometry (MS) analysis of SFK signaling. E. Phosphosite abundances under VEM (3 μM), DAS (50 nM), NAC (10 mM), and combinations thereof, measured by targeted MS. All treatments were delivered for 72 hours except NAC, which was delivered one hour before cell lysis. F. Immunoblot of phosphorylated SRC activation loop and total SRC under VEM (3 μM), NAC (5 mM, + and 10 mM, ++), and combinations thereof. G. Immunoblot of phosphorylated SRC activation loop and total SRC under H_2_O_2_ (30 μM) alone and in combination with NAC (10 mM). ^†^The tryptic phosphopeptide supporting SRC-pY419 also maps to activation loops on three other paralogous SFKs (FYN-pY420, LCK-pY394, and YES1-pY426).

We next subjected cells to direct measurement of intracellular ROS levels. In concordance with the observed transcriptional signature suggesting gradual oxidative stress, VEM-treated cells showed nearly 2.5-fold higher levels of ROS after 72 hours compared to control cells, and 1.5-fold higher levels compared to cells treated directly with 30 μM exogenous H_2_O_2_ (Figure 5C). ROS levels were restored to baseline or sub-baseline levels by one-hour treatment with N-acetylcysteine (NAC), an antioxidant prodrug and ROS scavenger. These results confirm that BRAF inhibition by VEM promotes significant yet sub-cytotoxic ROS accumulation.

To probe the relationship between VEM treatment, SFK signaling, and ROS accumulation, we treated cells with VEM, DAS, or the combination for 72 hours, with or without addition of NAC for one hour prior to harvesting, and subjected lysates to targeted MS, specifically aiming to quantify SFK phosphorylation and well-characterized SFK substrates (Figure 5D and Supplementary Table S4). We also quantified phosphorylated ERK1/2 to confirm successful VEM target engagement. The identity and quantitation of each phosphopeptide target was manually validated (Figures S5A and S5B). Targeted MS showed increased tyrosine phosphorylation on the SFK activation loop and on established SFK substrates under VEM treatment, which was reduced or fully prevented by addition of DAS (Figure 5E). Most substrates were downregulated by NAC treatment alone but to a lesser extent in combination with VEM, suggesting one hour of NAC treatment may not be sufficient time for PTPs to return aberrant phosphorylation to baseline levels.

We confirmed significantly increased SRC phosphorylation under VEM by western blot, which was prevented by combination treatment with DAS and with NAC in a dose-dependent manner (Figure 5F). Compared to MS-based quantitation, western blot showed NAC treatment had a greater effect on SRC phosphorylation, potentially due to ambiguity introduced by the non-unique mapping of the tryptic SFK phosphopeptide (LIEDNE(pY)TAR maps to human SRC, FYN, LCK, and YES1). Finally, we observed that direct exposure of cells to exogenous H_2_O_2_ drove an increase in SRC phosphorylation that was prevented by NAC (Figure 5G). Together, these results establish a causal association between VEM treatment, intracellular ROS levels, and net SFK activity.

### SFK-mediated adaptation to BRAF inhibition sensitizes cells to dasatinib

We sought to test whether early SFK induction by VEM would render cells susceptible to combination treatment with DAS. We observed pronounced and sustained cell growth inhibition under the drug combination across a panel of patient-derived BRAF^V600E^ melanoma cell lines, all of which eventually expanded under VEM alone (Figure 6A) and were unresponsive to DAS alone (Figure 6B). Two cell lines in particular, M233 and M381, were durably suppressed despite previous evidence of intrinsic resistance to VEM.^12^ These findings suggest some degree of generality of the SFK-mediated response to BRAF inhibition and confirm that this response can confer a strong mitogenic and survival dependency.

**Figure 6.**
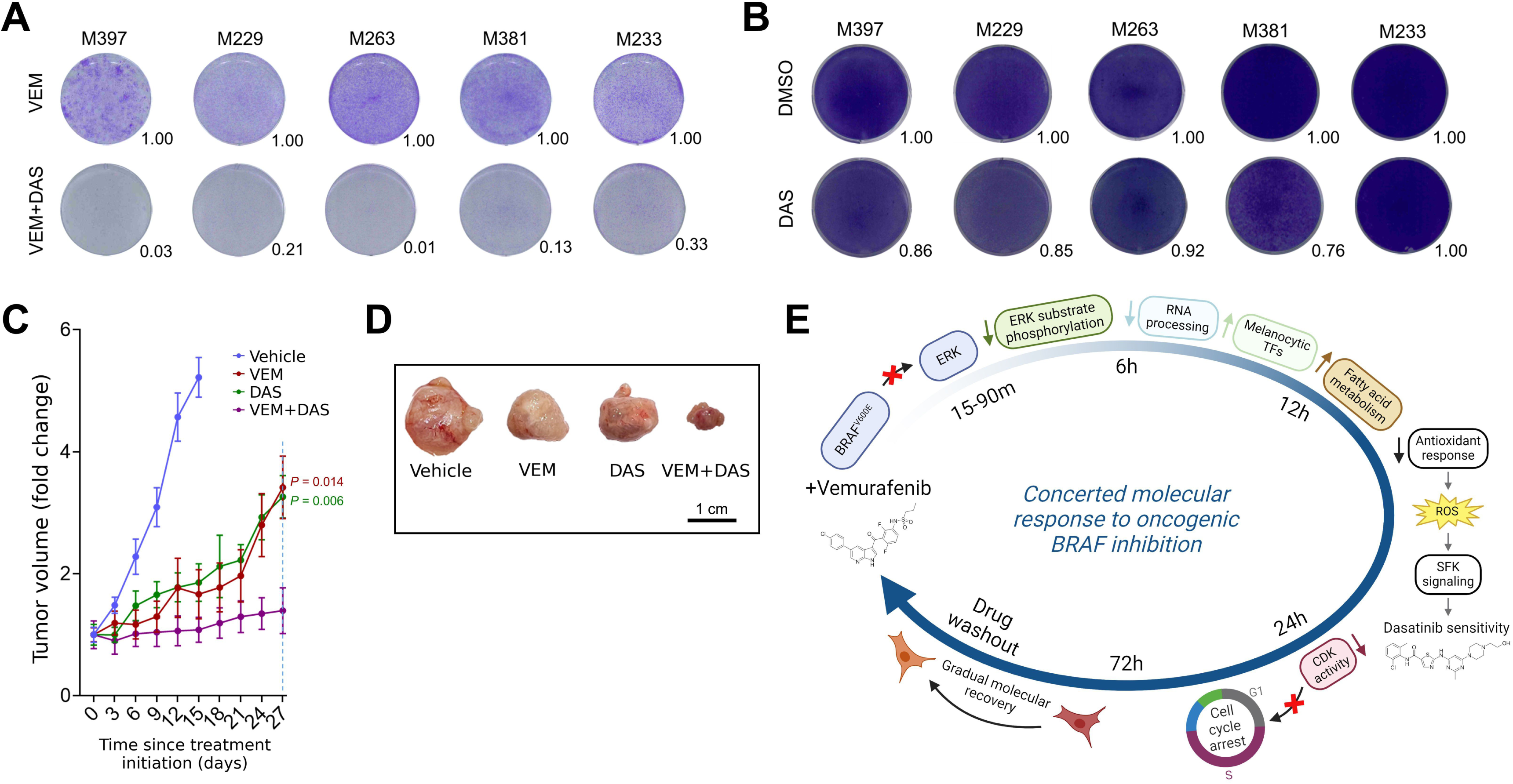
Adaptive response to BRAF inhibition confers sensitivity to SFK inhibitor dasatinib *in vitro* and *in vivo*. A. Clonogenic assays of a panel of patient-derived melanoma cell lines treated with VEM (3 μM) alone or in combination with DAS (50 nM). B. Clonogenic assays of a panel of patient-derived melanoma cell lines treated with DMSO (0.1%) or DAS (50 nM). C. Tumor growth curves of the M397-derived orthotopic xenograft mouse model treated with vehicle, VEM (100 mg/kg), DAS (30 mg/kg), or VEM combined with DAS. Error bars depict standard error of the mean across five replicates. P-values were calculated by two-sided Welch’s *t*-test. D. Resected tumors from cell line xenograft mouse models after 27 days of treatment with the indicated drugs. E. Timeline schematic of the dynamics of SFK activity and other selected critical regulatory events driven by BRAF inhibition by VEM.

We verified the efficacy of this combination in an orthotopic xenograft mouse model of the M397 line and observed significant tumor stasis with the drug combination compared to VEM and DAS monotherapy (Figure 6C and 6D). Notably, tumors were more responsive to DAS in vivo than the M397 line was in vitro; this is likely due to the higher dose of DAS delivered in vivo and also may indicate the presence of additional microenvironmental and physiological determinants of drug sensitivity that weren’t captured in simpler in vitro models.^31^ Taken together, our results support a time-resolved model where adaptation to BRAF inhibition leads to a gradual loss of redox homeostasis, accumulation of ROS, and resultant SFK activation, likely accompanied by PTP inhibition, and together leading to a pro-survival response that potently sensitizes cells to SFK inhibition (Figure 6E).

## Discussion

The systems-level intracellular molecular response to oncoprotein inhibition can dictate whether an exposed tumor cell will survive or die. In pursuing improved targeted therapies and therapy combinations for cancer, it has become crucial to map and better understand the complex and multifactorial systems that encode robustness to oncoprotein disruption. To elucidate the early temporal stages that precede and potentially lead to complete therapy resistance, we conducted a comprehensive analysis of the kinetic changes in cell signaling networks over minutes, hours, and days following the administration of vemurafenib (VEM), a BRAF^V600E^ inhibitor, using a melanoma cell line with partial responsiveness to BRAF inhibition as a model system. Given the importance of tumor cell-intrinsic signaling mechanisms in mediating responses to kinase inhibitors – including feedback inhibition, pathway cross-talk, and kinase redundancy, among others – we reasoned that a hypothesis-free time-resolved characterization of signaling events that unfold early under drug treatment would provide deep insights into cellular adaptation to targeted therapy.

We observed multiple distinct stages of signaling network remodeling in the drug-altered phosphoproteome, corresponding to early, intermediate, and late pathway alterations. Treatment with VEM lead to rapid and sustained inhibition of the BRAF-ERK axis and gradual downregulation of RB1 phosphorylation. These observations indicate reduced mitotic activity and were consistent with corresponding dynamic measurements of cell viability. We observed distinct kinetics of phosphorylation loss on canonical ERK1/2 substrates in response to VEM treatment, including ERF, MAP2K2 (MEK2), and STMN1, suggesting that their phospho-regulation may be controlled by other kinases besides ERK1/2 or by distinct serine/threonine phosphatases.

The marked suppression of ERK1/2 activity during VEM treatment suggests that cell survival was not dependent on reactivation of BRAF or ERK1/2 during the measured time course. We speculate that the absence of ERK1/2 reactivation within our model system may be attributed to the efficient blockade of both mutant and wild-type BRAF by VEM at 3 μM, such that the RAF-ERK pathway is durably inhibited even in the presence of possible upstream RTK/RAS activation.

Through the kinetic profiling of mRNA transcript levels, we found strong agreement in cell state dynamics with the time-resolved phosphoproteome. Both classes of biomolecules demonstrated an early response to VEM and a near-complete reversal of remodeling following six days of VEM removal. These melanoma cells are known to exhibit significant molecular and phenotypic plasticity under BRAF inhibition;^12^ our results confirm that maintenance of the early drug-adapted cell state depends on sustained BRAF inhibition.

Inference of kinetically-defined signaling and transcriptional modules recovered known associations while also revealing many examples of new signaling-gene regulatory associations. Although it is currently intractable to individually examine and experimentally test the relationships between all possible pairs of observed phosphosites and genes (>17M), we believe this time-resolved multimodal dataset serves as a valuable resource to investigate new lines of inquiry into the concerted molecular processes that follow oncoprotein inhibition and tumor cell stress within a well-characterized model system of adaptive drug response.

SFK induction during targeted therapy treatment has been described in multiple cancers, including melanoma;^30,31,32^ our study sheds light on the kinetics associated with this response, revealing that signs of SFK induction can emerge as early as 3-6 hours of oncoprotein inhibition. This onset is markedly earlier than previously appreciated, suggesting a more immediate cellular adaptation to targeted therapy via this axis than previously appreciated. Intriguingly, similar to the distinct phosphorylation patterns of different canonical ERK1/2 substrates, our analysis revealed distinct patterns of phosphorylation on canonical SFK substrates – for instance, after 72 hours PXN-pY118 was upregulated 1.8-fold, CTTN-pY421 was upregulated 4.1-fold, and TNS1-pY1427 was upregulated over 5.2-fold on average compared to control cells. This variation in signaling magnitude may suggest the engagement of distinct sets of tyrosine phosphatases and kinases, including but not limited to SFKs, in modulating the phosphorylation states of these crucial substrates.

We observed a substantial increase in intracellular ROS levels following oncogenic BRAF inhibition. By perturbing cells with ROS with or without NAC as a competitive antioxidant, we were able to isolate the effect of ROS on SFK activity and found that increased ROS level leads to increased SFK activation loop phosphorylation. In addition to their established role in directly activating SFKs by oxidizing catalytic cysteine residues,^34,35,36^ ROS are known to modulate signaling networks by inhibiting the action of PTPs, similarly by oxidizing the free thiol groups on catalytically essential cysteines.^37,38,44^ While our study did not directly identify PTP oxidation as an explanatory mechanism for the net increase in tyrosine phosphorylation under VEM, it is likely that PTP inhibition contributed to this observation. We note that while acute treatment of NAC was sufficient to reduce SFK phosphorylation after VEM exposure, this effect did not extend to many canonical SFK substrates when we profiled by targeted mass spectrometry (MS), perhaps indicating that one hour of NAC exposure is not enough time to fully recover the catalytic activity of PTPs.

Our observations reveal a strong degree of synergy between VEM and the SFK inhibitor dasatinib (DAS) in vitro, and we observed tumor stasis with the combination in vivo despite inability of either monotherapy to control tumor burden. The therapeutic potential of DAS as a monotherapy for melanoma has been explored in multiple clinical trials, which have generally failed to demonstrate a clinically meaningful response within a tolerable dose range.^45,46^ DAS is currently approved only for the treatment of chronic myeloid leukemia and a subset of other Philadelphia chromosome-positive leukemias due to its potent activity against BCR-ABL1. Consistent with the expectation of low sensitivity to SFK inhibition alone, we observed little to no effect of DAS across a panel of five patient-derived melanoma cell lines. However, the potent and sustained inhibition of cell growth when combined with VEM suggests that induction of SFK signaling by VEM reveals an underlying synthetic lethal interaction between mutant BRAF and the SRC family, which may be exploited therapeutically. Additional preclinical studies would be needed to verify the generality of this drug combination’s efficacy and to identify potential biomarkers identifying the subset of patients most likely to benefit from low-dose SFK inhibition combined with BRAF/MEK blockade.

This study underscores the significant potential of quantitatively characterizing the dynamics of early alterations in cell signaling and transcriptional networks under targeted therapy treatment. Such analysis provides powerful insights into multiple aspects of cell state: the dynamics of drug-target engagement, the downstream molecular processes regulated by the target, and the induction of compensatory pro-survival signaling pathways that should be considered for co-targeting.

## Acknowledgements

We thank Antoni Ribas for providing the patient-derived melanoma cell lines. This work was funded by NIH grants U01 CA217655 and U54 CA274509, and supported by the MIT Center for Precision Cancer Medicine. W.W. was funded by the Andy Hill CARE Fund. C.T.F. was funded by a graduate fellowship from the MIT Ludwig Center.

## Author contributions

J.R.H., W.W., and F.M.W. conceived and supervised the study. C.T.F., C.L., W.W., and F.M.W. designed the experiments. C.T.F., C.L., H-Y.C., X.Y., and H.C. performed the experiments. C.T.F., C.L., H-Y.C., J.R.H., W.W., and F.M.W. analyzed the data. The manuscript was written by C.T.F., C.L., J.R.H., W.W., and F.M.W., with feedback from all other authors.

## Data and code availability

The processed time-series phosphoproteomics and RNA-seq data are provided in Supplementary Tables S1 and S2, respectively. The results of the regulatory module inference are provided in Supplementary Table S3. The processed targeted phosphoproteomics data are provided in Supplementary Table S4. The raw and searched MS phosphoproteomics data have been deposited to the ProteomeXchange Consortium via the PRIDE partner repository with the dataset identifier PXD048479. The raw fastq-formatted files as well as processed RNA-seq data are available on NCBI GEO via accession number GSE252781. The python code for reproducing the signaling and transcriptional module inference is available at the following GitHub repository: github.com/flowerc/ModuleInference.

## Materials and methods

### Cell lines and drug treatment

M-series patient-derived cell lines were generated under UCLA institutional review board approval # 11–003254 and transferred to Institute for Systems Biology under established materials transfer agreement (MTA). Cells were cultured in a water-saturated incubator at 37 °C with 5% CO_2_ in RPMI-1640 containing L-glutamine (Gibco, catalog # 11875093) supplemented with 10% fetal bovine serum (ATCC) and 0.2% MycoZap Plus-CL antibiotics (Lonza, catalog # 195261). Cell lines were routinely tested for mycoplasma and were periodically authenticated to its early passage using the GenePrint 10 System (Promega, catalog # B9510). VEM and DAS (Selleck Chemicals, catalog #s S1267 and 1021, respectively) were dissolved in DMSO at designated concentrations before applying to cell culture media at a final DMSO concentration of 0.1% (v/v). Control cells were treated with 0.1% DMSO. NAC (Sigma-Aldrich, catalog # A7250) was dissolved in phosphate-buffered saline. For phosphoproteomics and RNA-seq experiments, M397 cells were grown in 10 cm tissue culture plates, plated at 60% confluency, and treated for the specified amount of time before being harvested from culture.

### Cell viability dose-response assay

M397 cells were seeded in a 96-well plate (3000 cells/well) in triplicate and incubated for 24 hours, followed by treatment with VEM in 0.1% DMSO at the designated doses for 72 hours. Cell viability was then quantified using CellTiter-Glo (Promega, catalog # G7572) according to the manufacturer protocol. All luminescence values were normalized to the average of the three drug-free control wells. Conventional monophasic dose-response models achieved poor fit to the data due to the clear biphasic nature of the M397 response to VEM; therefore, the following modified viability model was derived to accommodate the biphasic dose response:

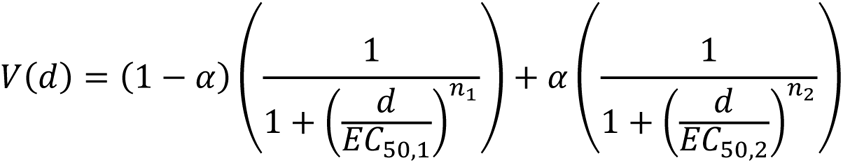

where is the fraction of viable cells under dose (in nM). Under this model specification, once fitted to the data, and are estimates of the doses at which there is a half-maximal effect for the first and second dose-response phases, respectively. Similarly, and are measures of the steepness of the first and second dose-response phases, respectively. provides an estimate of the intermediate viability of cells treated with doses between and . Model fitting was performed using the curve_fit function in the SciPy scientific computing python package.^47^

### Dead cell quantitation and time-resolved viability assay

M397 cells were seeded in 6 cm plates (0.2M/plate), incubated for 24 hours, and treated with 0, 3, or 100 μM VEM in 0.1% DMSO for 72 hours. Six biological replicates were seeded per condition. 100 μM VEM was used as a positive control for cell death. Media was collected from each plate, and cells adhered to the plate were trypsinized and added to the collected media. Cells were pelleted, resuspended in PBS, and stained with trypan blue. Dead (trypan blue-positive) and live (negative) cells were quantified using a hemocytometer. For quantifying cell viability kinetics, M397 cells were plated onto six-well plates (0.1M/well), incubated for 24 hours, and treated with 3 μM VEM or 0.1% DMSO. Three biological replicates were seeded per condition. Cell viability was monitored by an Incucyte S3 Live-Cell Analysis System (Sartorius). Phase contrast images were taken every six hours throughout a 72-hour period. Cell numbers were quantified using Incucyte classic confluence analysis workflow.

### Phosphoproteomics sample preparation

M397 cells were lysed in 8M urea and total protein was quantified by bicinchoninic acid assay (Pierce, catalog # 23225). For each sample, 700 μg of protein was subjected to reduction in 10 mM dithiothreitol at 56°C for one hour, followed by cysteine alkylation in 55 mM iodoacetamide at room temperature for one hour in the dark. Samples were diluted 5X in 100 mM ammonium acetate to decrease the urea concentration prior to protein digestion, then samples were incubated with sequencing-grade modified trypsin (Promega, catalog # V5113) for 18 hours at a trypsin:protein ratio of 1:50 (w/w). Digestion was halted with 10% acetic acid, and samples were desalted by solid phase extraction using Sep-Pak C18 Plus Light cartridges (Waters, catalog # WAT023501) and eluted in 40% acetonitrile. All samples were concentrated in a vacuum centrifuge. Peptide concentration was quantified by bicinchoninic acid assay (Pierce), and 120 μg of peptide from each sample was freeze-dried by lyophilization. Samples were resuspended in 50 mM HEPES and labeled with TMTpro isobaric mass tags (Thermo Fisher Scientific, catalog # A52045) at a TMT:peptide ratio of 4:1 (w/w) for five hours at room temperature. Unreacted TMT was quenched with 3.2 μL of 5% hydroxylamine, then samples within each plex were pooled and dried by vacuum centrifugation.

### Phospho-motif immunoprecipitation and phosphopeptide enrichment

TMT-labeled samples were resuspended in immunoprecipitation (IP) buffer (100 mM Tris-HCl, 1% NP-40, pH 7.4). For IP of tyrosine phosphorylated peptides, samples were incubated with 24 μg of anti-phosphotyrosine 4G10 antibody (BioXCell, catalog # BE0194) and 12 μg of anti-phosphotyrosine PT-66 antibody (Sigma-Aldrich, catalog # P3300) conjugated to protein G agarose beads (Sigma-Aldrich, catalog # IP04). For IP of phosphopeptides containing the (pS/pT)Q and (pS/pT)P phospho-motifs, supernatant from the phosphotyrosine IP was incubated with 40 μL of anti-(pS/pT)Q and anti-(pS/pT)P antibody pre-conjugated to agarose beads (Cell Signaling Technologies, catalog #s 12267 and 14378S, respectively). Immunoprecipitation was carried out over 18-24 hours at 4°C. Peptides were eluted with 0.2% trifluoroacetic acid for 10 minutes and subjected to an additional phosphopeptide enrichment step using High-Select™ Fe-NTA spin columns (Thermo Fisher Scientific, catalog # A32992) to reduce the abundance of non-phosphorylated peptides which may have bound nonspecifically to the IP beads. For phosphoserine/threonine enrichment (with no phospho-motif enrichment), 2 μL of peptide from the IP supernatant was diluted in 50 μL of 0.2% trifluoroacetic acid and transferred to an Fe-NTA spin column. All phosphopeptide samples were eluted according to the manufacturer’s instructions, dried by vacuum centrifugation, and resuspended in 5% acetonitrile in 0.1% formic acid.

### Liquid chromatography and mass spectrometry

Each phosphopeptide mixture corresponding to a single TMT plex was loaded directly onto a chromatography column using a helium packing device in order to minimize sample loss. Columns were constructed in-house; fused silica capillary with inner diameter 50 μm and outer diameter 200 μm (Molex, catalog # 1068150017) was cut to a length of 25 cm and pulled using a micropipette laser puller to create an integrated emitter tip with 1-2 μm inner diameter, then was packed with 10 cm of 5 μm C18 beads (YMC Co, catalog # AQ12S05) and conditioned and tested using a tryptic digest of bovine serum albumin. Liquid chromatography-tandem mass spectrometry was performed using an Agilent (1100-Series) chromatograph coupled to a Thermo Scientific™ Orbitrap Exploris™ 480 mass spectrometer. Peptides were separated using 0.2M acetic acid (solvent A) and 70% acetonitrile in 0.2M acetic acid (solvent B) over the following 140-minute gradient profile: (min:%B) 0:0, 10:11, 115:32, 125:60, 130:100, 133:100, 140:0. Electrospray ionization was carried out at 2.5 kV and at a flow rate of approximately 100 nL/min (200 μL/min through a flow splitter achieving a ∼1:2000 split). For the time-resolved phosphoproteomics experiment, the mass spectrometer was operated in data-dependent acquisition mode as follows: in each cycle, a full MS scan of precursor ions with m/z between 380-2000 was acquired with a resolution of 60,000 at 200 m/z and AGC target of 3e6, followed by a series of MS/MS scans for a total cycle duration of 3 s. For MS/MS scans, precursors with charge state between 2-6 and intensity greater than 1e4 were isolated with an isolation width of 0.4 Th and accumulated until the AGC target of 1e5 was reached, or until 250 ms had elapsed. Fragmentation was performed using a normalized collision energy of 33%, and fragment ions were scanned with a resolution of 60,000 at 200 m/z. Each precursor was allowed to be selected for MS/MS twice before being dynamically excluded for 45 s. For the targeted phosphoproteomics experiment, the mass spectrometer was restricted to selecting ions for MS/MS based on an inclusion list of mass-to-charge ratios of peptides of interest from the time-resolved phosphoproteomics data (Supplementary Table S4). The mass tolerance for the inclusion list was set to 3 parts per million (ppm). Given the relatively limited set of monitored phosphopeptide targets, the acquisition sensitivity was boosted by increasing the resolution (full MS and MS/MS) to 120,000, increasing the AGC target for MS/MS to 1e6, and disabling the intensity threshold and dynamic exclusion.

### Phosphopeptide identification and quantitation

Raw files containing phosphopeptide mass spectra were processed using Proteome Discoverer version 3.0 (Thermo Fisher Scientific) and searched using Mascot version 2.4 (Matrix Science). For the time-resolved phosphoproteome experiment, spectra were searched against the canonical human proteome (SwissProt reviewed sequences, version 2023_01) with trypsin digestion allowing for up to one missed cleavage, and with a precursor ion mass tolerance of 5 ppm. For the targeted phosphoproteomics experiment, spectra were searched against a custom database containing the phosphopeptide sequences that were specifically targeted for acquisition, with no enzymatic digestion, and with a precursor ion mass tolerance of 3 ppm. For both phosphoproteomics experiments, MS/MS spectra were searched with a fragment ion mass tolerance of 20 mmu. Precursor ions and TMT reporter ions were removed from MS/MS spectra prior to searching using the non-fragment filter node in Proteome Discoverer. Cysteine carbamidomethylation, TMT-labeled lysine, and TMT-labeled peptide N-termini were set as static modifications; phosphorylation of tyrosine, serine, and threonine, and oxidation of methionine were set as dynamic modifications. Phosphorylation site localization was performed using ptmRS^48^ within Proteome Discoverer. For peptide quantitation, TMT reporter ion intensities were extracted with an integration tolerance of 10 ppm and were isotope-corrected using Proteome Discoverer according to the manufacturer-provided batch-specific isotopic impurities of each TMT channel.

Peptide-spectrum matches (PSM) were exported from Proteome Discoverer and filtered by match quality (expectation value < 0.05) and phosphosite localization confidence (probability > 0.9 for all phosphosites on the peptide). Any PSM corresponding to a spectrum that matched to more than one amino acid sequence with equal expectation value was removed. PSMs with more than four missing TMT values were discarded. For the remaining PSMs, missing TMT reporter ion intensities were replaced with a value equivalent to half the intensity of the least-intense fragment ion observed in the corresponding MS/MS spectrum; this approach assumes that an absent reporter ion would have been detected and recorded into the raw file had its intensity been greater than the least-intense fragment in the spectrum. Under this assumption, the intensity of the absent reporter lies between zero (“true absence”) and the intensity of the minimal fragment, with uniform probability. TMT reporter ion intensities were then summed across PSMs sharing a common modified peptide sequence. Phosphopeptides derived from the same source protein(s) with shared phosphorylation site pattern (for example, two phosphopeptides covering the same phosphosite but with different tryptic cleavage pattern) were aggregated to eliminate phosphosite redundancy in the resulting data matrix.

### RNA-seq

M397 cell pellets were snap frozen at the designated timepoints after treatment with 3 μM VEM or 0.1% DMSO. Total RNA was extracted from cell pellets using AllPrep DNA/RNA kit (Qiagen, catalog # 80204). The mRNA library was constructed with TruSeq RNA Sample Preparation Kit (Illumina, catalog # RS-122-2001) and sequenced on an Illumina HiSeq 2500 System with paired-end 150-base-pair reads following the manufacturer’s recommendations by Novogene.

### Immunoblotting

Antibodies against phospho-SRC (Y416), total SRC, and GAPDH (Cell Signaling Technology, catalog #s 2101S, 2109T, and 5174S, respectively) were used. The Invitrogen precast gel system NuPAGE was used for running the western blot. The 4–12% Bis-Tris gels were loaded with samples. After blotting, the membranes were blocked in 5% BSA with TBS + 0.1% Tween-20 (TBST) mix for at least one hour at room temperature. Membranes were then incubated overnight with the primary antibody in 5% BSA with TBST at 4°C. The next day, membranes were washed three times for five minutes in TBST, incubated with a suitable HRP-coupled secondary antibody for one hour at room temperature, and washed three times. Proteins were visualized with SuperSignal™ West Pico PLUS Chemiluminescent Substrate (Thermo Fisher Scientific, catalog # 34577) using the ChemiDoc™ XRS+ System.

### Clonogenic assay

Cells were seeded onto six-well plates at an optimal seeding density determined from each cell line’s specific growth rate. For the VEM plus DAS experiment (Figure 6A), cells were treated with VEM monotherapy or VEM plus 50 nM DAS for the same amount of time. The dose of VEM used was 2-fold the cell line-specific EC50 which was previously determined.^12^ The media (with drug or DMSO) was replenished every two days. For the combinatorial treatment, the treatment was stopped followed by staining when the VEM-treated cells displayed clear regrowth. For the toxicity assessment (Figure 6B), cells were treated with 50 nM DAS or DMSO and the treatment was stopped followed by staining after five days. Upon the time of staining, 4% paraformaldehyde was applied onto colonies to fix the cells and 0.05% crystal violet solution was used for staining the colonies. ColonyArea, an ImageJ plugin, was used to perform standard analysis of colony formation assays.

### ROS production assay

M397 cells were split into six groups and treated as follows: 0.1% DMSO; 3 μM VEM; 3 μM VEM plus 10 mM NAC; 30 μM H_2_O_2_; 30 μM H_2_O_2_ plus 10 mM NAC; 0.1% DMSO plus 10mM NAC. All treatments were given for three days except NAC, which was given one hour prior to ROS quantitation. ROS levels were quantified by ROS-Glo™ H_2_O_2_ Assay (Promega, catalog # G8820) according to the manufacturer’s protocol. The luminescence signals were measured by the Synergy H4 Plate Reader.

### Melanoma mouse xenograft experiments

Mice were maintained in a specific pathogen-free (SPF), temperature-controlled (21-23°C) animal facility on a reverse 12-hour light, 12-hour dark cycle at the University of Washington. Food and water were given ad libitum. Mice were kept under animal bio-safety level (ABSL-2) conditions and handled under protocols approved by the Institutional Animal Care and Use Committee (IACUC) of the University of Washington (PROTO201900025) in accordance with international guidelines. To generate xenograft mouse tumor models, 6 to 7-week-old NOD. Cg-Prkdc^scid^ Il2rg^tm1Wjl^/SzJ (NSG) female mice were purchased from the Jackson Laboratory (catalog # 005557). NSG mice were inoculated with M397 cells (10^6^ cells per mouse) in Matrigel mixture (Corning, catalog # 354234) via a subcutaneous injection into the right flank. When the average tumor volume reached ∼200 mm^3^, the mice were randomized into four groups: control, 100 mg/kg VEM, 30 mg/kg DAS, and 100 mg/kg VEM plus 30 mg/kg DAS, with five mice per group. VEM and DAS were formulated as 5% DMSO in 0.5% methyl cellulose and given once daily by oral gavage. Tumor length and width were measured once every three days using calipers. Tumor volume was calculated using the equation Volume (mm^3^) = Length (mm) × Width (mm^2^) / 2. Mice were euthanized and tumors were excised at day 27, except for the mice in the control group that were sacrificed earlier when the average tumor volume reached 1500 mm^3^.

### Statistical analysis software

Statistical testing, principal components analysis, and hierarchical clustering of the phosphoproteomics data was performed using the SciPy, scikit-learn, and seaborn python packages. For principal components analysis (Figure 1E) and pairwise sample correlation analysis (Figure S1C), phosphopeptide abundance values were mean-normalized.

### RNA-seq analysis

Reads were aligned against the human genome (hg19) using STAR (v2.7.3a). Read counts were quantified using htseq-count^49^ with known gene annotations from UCSC.^50^ Transcript level was calculated as fragments per kilobase of transcript per million mapped reads (FPKM). Transcript levels (FPKM+1) were centralized and scaled prior to principal components analysis (Figure 2A) using scikit-learn. Gene-set enrichment analysis (GSEA) was conducted using GSEA v4.1.0 software with 1000 permutations and weighted enrichment statistics. Normalized enrichment score was computed across the curated Molecular Signatures Database (MSigDB) Hallmark, C2, and C5 GO:BP gene sets.

### Transcription factory regulatory network analysis

The NetAct^51^ R package was used for constructing the TF regulatory network using time-series transcriptome data, depicted in Figure S2. Specifically, each RNA-seq sample was randomly downsampled to 3 replicates by seqtk. Each replicate recapitulates 50% of the original RNA-seq reads. Read counts quantified by htseq-count were used as the input for NetAct to construct core TF regulatory networks using inferred gene activities. NetAct uniquely integrates both generic TF-target relationships from literature-based databases and context-specific gene expression data to construct an optimized TF-target gene set database. NetAct takes the following steps to construct TF networks from raw read counts: (1) A pre-processing function called preprocess_counts() was used to filter out lowly expressed genes and retrieve associated gene symbols for raw count data. (2) Pairwise comparisons were defined to extract differentially expressed genes at each time point over the 3-day drug treatment procedure, resulting in a total of 15 group comparisons across six time points: control, and post-drug treatment at 12h, 24h, 36h, 48h, and 72h. Differentially expressed genes are subsequently identified using the limma package^52^ with q-value ≤ 0.05. The resulting normalized expression data is saved alongside the phenoData metadata into the standard ExpressionSet object for downstream analysis. (3) A permutation approach was adopted to select TFs with significantly altered activities across the drug treatment by using the gene set enrichment analysis against literature-based TF-target databases. The TFs are aggregated from the multiple comparisons with q-value ≤0.01. The activities of enriched TFs were further calculated from the standardized expression levels of their target genes with weighting factors defined as a Hill function. (4) The TF regulatory networks were constructed by using both the TF-TF regulatory interactions from the TF-target database and the activity values. The link interactions were filtered out based on mutual information and entropy. The plot_network() function was used to interact with the network topology.

### Regulatory module inference

The signaling and transcriptional module inference depicted in Figure 3 was performed as follows: TMT intensity values were averaged across the three replicates for each phosphosite in the time-series phosphoproteomic data, and the DMSO control samples were dropped. Data were z-score normalized (phosphosite-wise) and a pairwise Pearson correlation matrix was computed. Euclidean distances were calculated between each pair of phosphosites and used for hierarchical clustering, which was applied to the correlation matrix. A threshold on the cophenetic distance proportional to the total number of phosphosites was applied to the dendrograms, resulting in discrete clusters of phosphosites. The gene IDs within each cluster were queried for over-and under-representation of gene sets using Fisher’s exact test and the following libraries of gene sets downloaded from MSigDB (all version 2023_1): BioCarta, GObp, GOcc, GOmf, Hallmarks, KEGG, PID, Reactome, TFT, WikiPathways. P-values were corrected for multiple hypotheses by the Benjamini-Hochberg procedure. These steps were also applied to the RNA-seq data, after transcripts where quantitation (FPKM) was 0 in any sample were filtered out. For each module, the consensus dynamics were calculated as the mean z-score-normalized abundance across all members of the module. Consensus dynamics were then used to calculate meta-correlations between every pair of inferred signaling and transcriptional modules, as depicted in Figure 3C. P-values for meta-correlation were determined based on how frequently two randomly-sampled multivariate normally-distributed random variables achieve a Pearson correlation coefficient at least as extreme as the observed meta-correlation, followed by multiple hypothesis correction by the Benjamini-Hochberg procedure.

### Data visualization

Results were visualized using the following python packages: matplotlib, seaborn, holoviews. The schematics depicted in Figures 1C, 5D, and 6E were created using BioRender.

---

**Figure S1.**
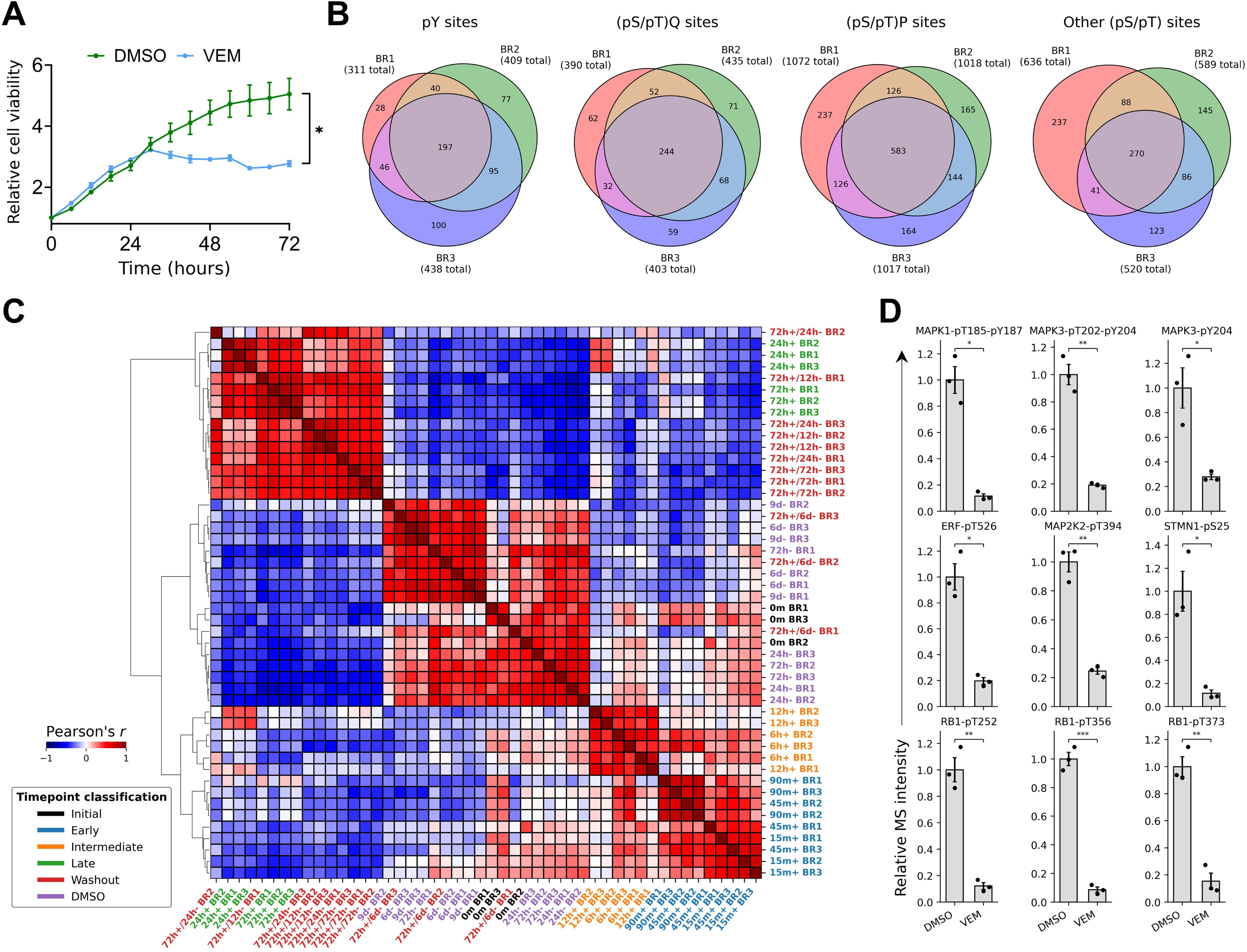
Growth and signaling characteristics of M397 melanoma cells under BRAF inhibition, related to Figure 1. A. Viability of melanoma cells under 0.1% DMSO or 3 μM VEM over a three-day time course. Error bars depict standard error across three replicates. * : *P* < 0.05 by two-sided Welch’s *t*-test. B. Number of high-confidence phosphosite identifications by phosphoproteomics. Values are reported for each biological replicate, each of which constitutes a single multiplexed MS analysis, and are segregated by each serially-enriched phospho-motif. C. Pairwise sample correlation and hierarchical clustering derived from phosphoproteomic data. D. Abundances of phosphosites on MAPK1 and MAPK3 (ERK2 and ERK1, respectively) (top), canonical ERK1/2 substrates (middle), and RB1 (bottom) at 72 hours under VEM or DMSO. Error bars depict standard error of the mean across three replicates. * : *P* < 0.05; ** : *P* < 0.01; *** : *P* < 0.005; by two-sided *t*-test with multiple hypothesis correction by the Benjamini-Hochberg procedure.

**Figure S2.**
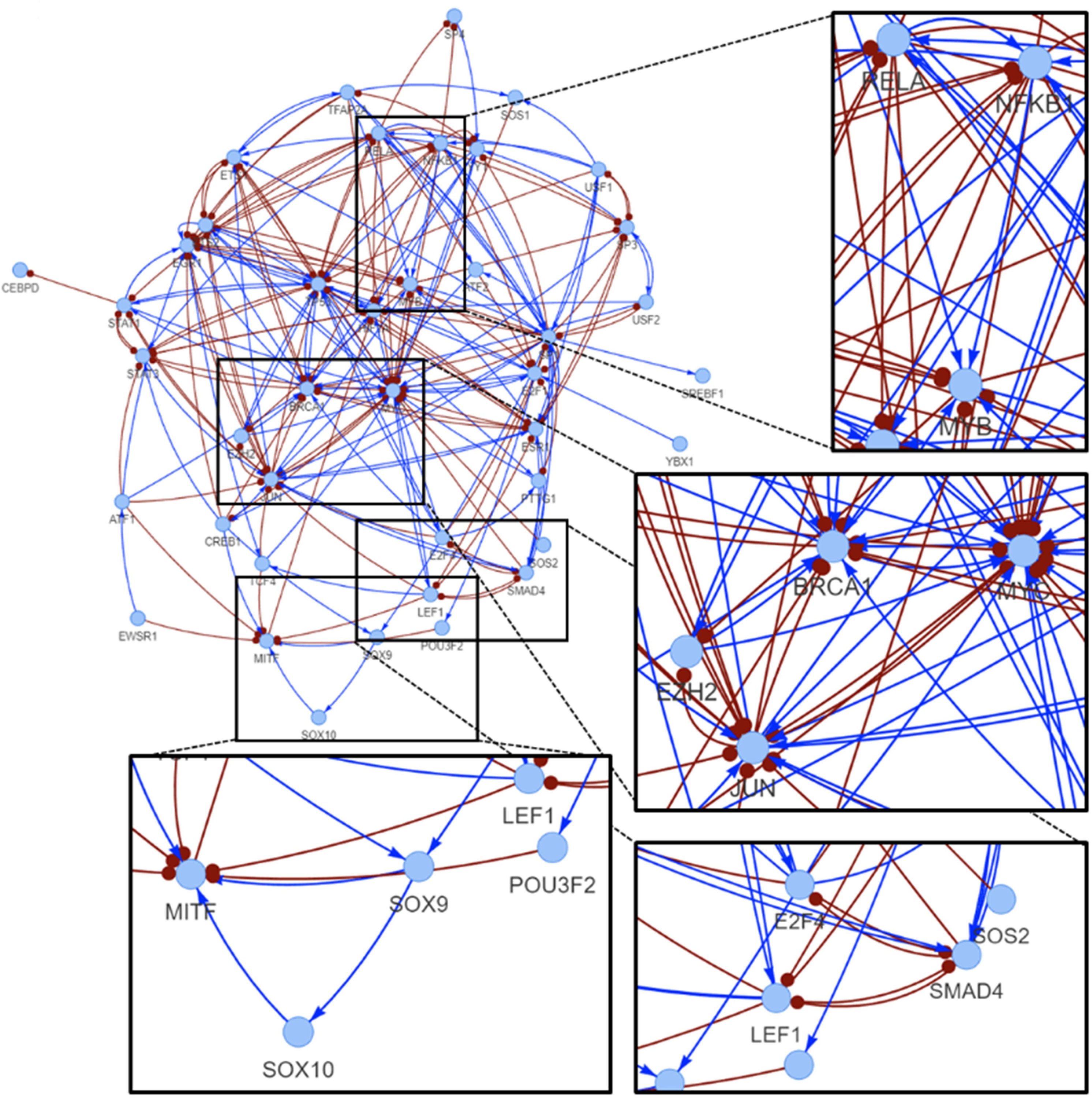
Remodeling of gene regulation by VEM, related to Figure 2. Inferred transcription factor regulatory network under VEM treatment for 72 hours, with representative zoomed-in sections. Blue lines and arrowheads signify gene activation; red lines and blunt heads signify gene inhibition.

**Figure S3.**
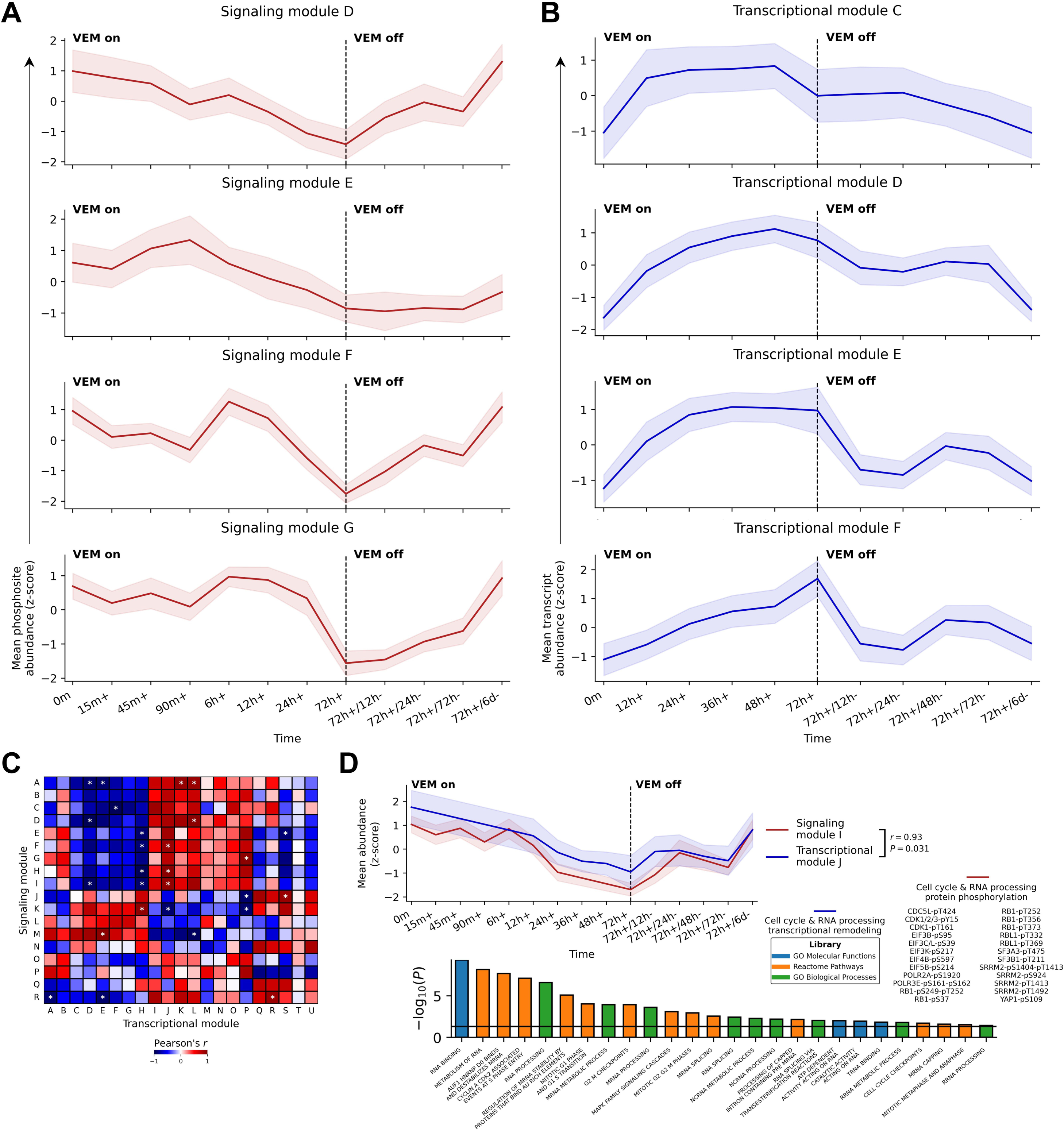
Integrative analysis of signaling and transcriptional response to BRAF inhibition, related to Figure 3. A. Consensus dynamics of qualitatively similar but kinetically distinct signaling modules D-G under VEM. Shaded regions depict standard deviation across all members of the depicted module. B. Consensus dynamics of qualitatively similar but kinetically distinct transcription modules C-F under VEM. Shaded regions depict standard deviation across all members of the depicted module. C. Pairwise correlation of signaling and transcription modules. * : *P* < 0.05. *P*-values indicate the estimated probability of uncorrelated modules producing an *r* value at least as extreme as the observed value, with multiple hypothesis correction by the Benjamini-Hochberg procedure. D. Consensus dynamics of signaling module A and transcription module E under VEM (top left); phosphosites related to ERK1/2 pathway signaling in signaling module A (top right); overrepresented gene sets in transcription module E related to metabolic rewiring and fatty acid oxidation (bottom). Shaded regions depict standard deviation across all members of the depicted module. *P*-values were derived from two-sided Fisher’s exact test with multiple hypothesis correction by the Benjamini-Hochberg procedure.

**Figure S4.**
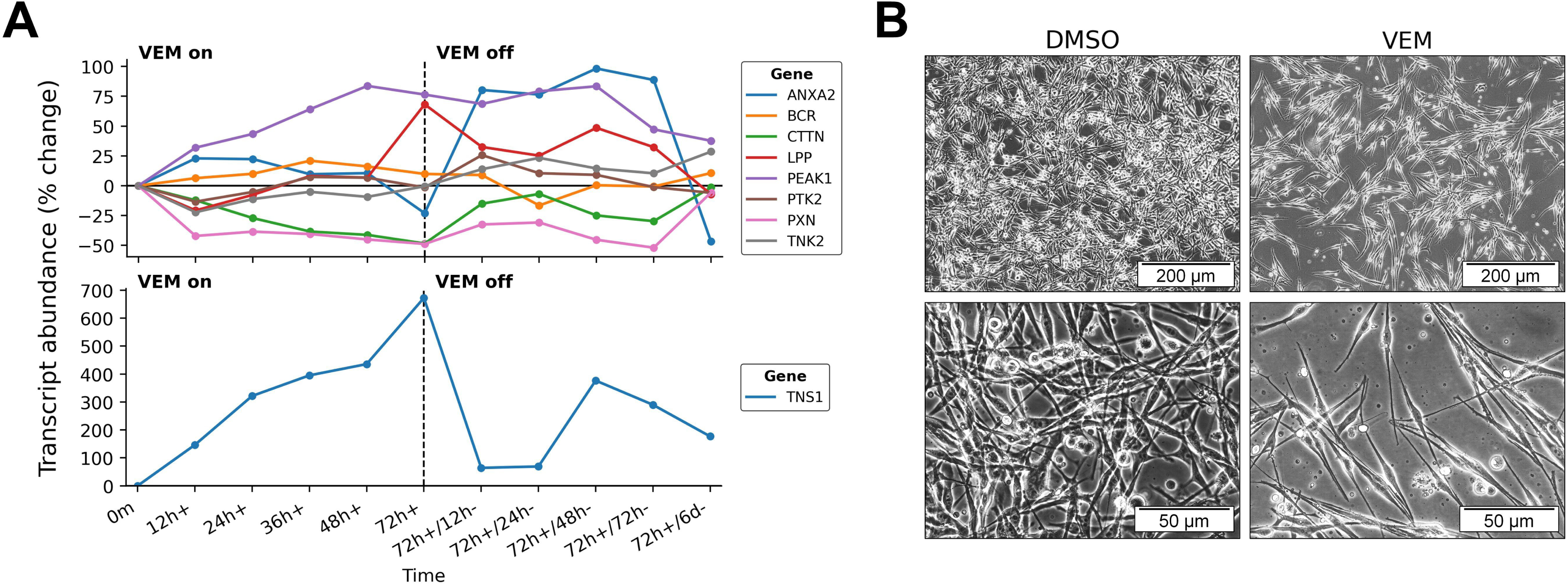
Transcriptional dynamics of SFK substrates and morphological characteristics under BRAF inhibition, related to Figure 4. A. mRNA dynamics of SFK substrate genes shown in Figure 4D. B. Morphology of M397 cells under DMSO or VEM for 72 hours.

**Figure S5.**
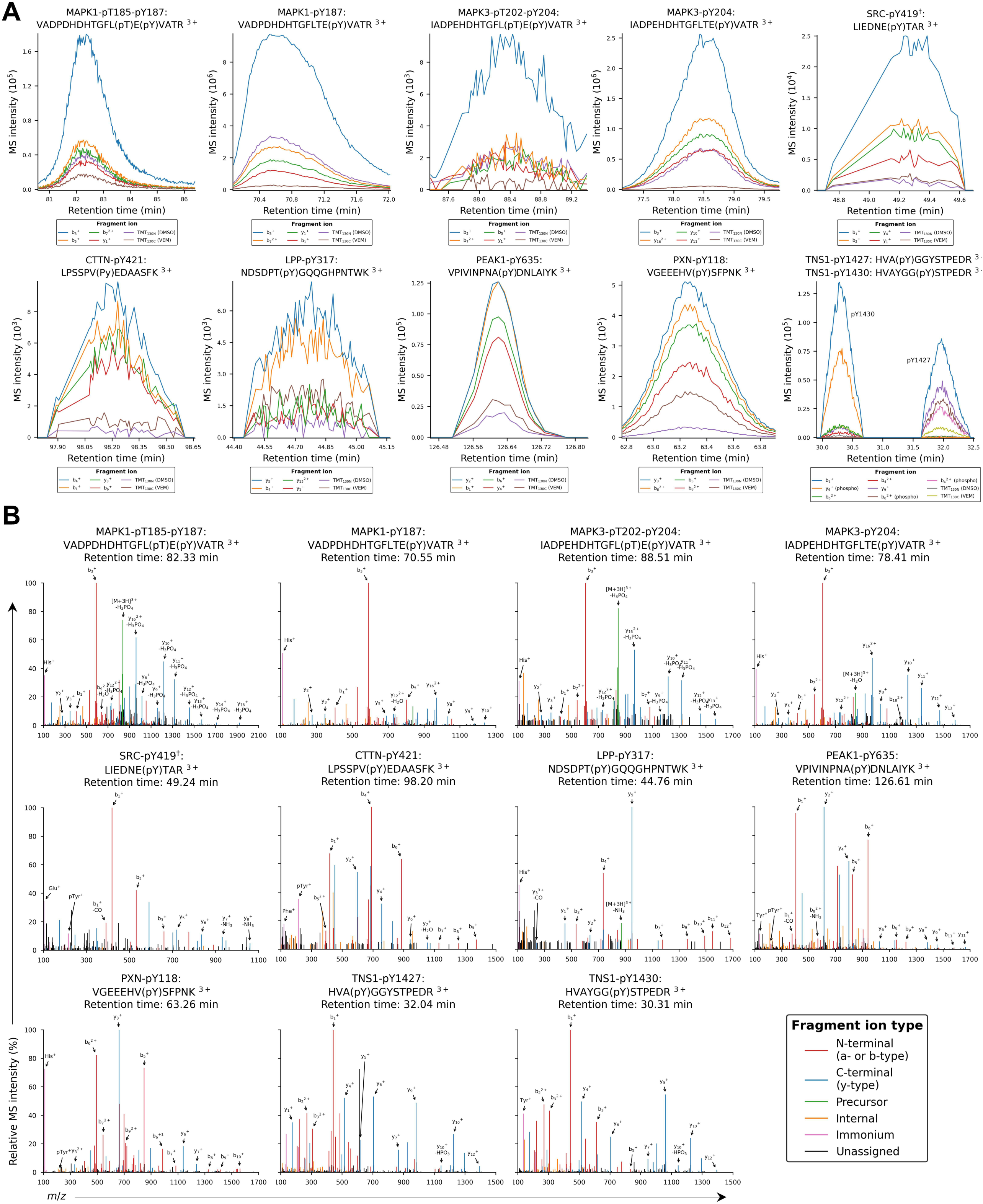
Manually validated targeted MS data, related to Figure 5. A. Fragment ion chromatograms of the phosphopeptides quantified in Figure 5E. B. MS/MS spectra of the phosphopeptides quantified in Figure 5E, acquired at the elution apex where sensitivity is maximal. Tandem mass tag reporter ions used for quantitation are not shown. ^†^The tryptic phosphopeptide supporting SRC-pY419 also maps to activation loops on three other paralogous SFKs (FYN-pY420, LCK-pY394, and YES1-pY426).

